# Mortality in trauma patients admitted during, before, and after national academic emergency medicine and trauma surgery meeting dates in Japan

**DOI:** 10.1101/453233

**Authors:** Tetsuya Yumoto, Hiromichi Naito, Hiromi Ihoriya, Takashi Yorifuji, Atsunori Nakao

**Affiliations:** Department of Emergency, Critical Care, and Disaster Medicine, Okayama University Graduate School of Medicine, Dentistry and Pharmaceutical Sciences, Okayama, Japan; Department of Human Ecology, Okayama University Graduate School of Environmental and Life Science, Okayama, Japan

**Author notes:** Corresponding author: (TYu).

## Abstract

Annually, many physicians attend national academic meetings. While participating in these meetings can have a positive impact on daily medical practice, attendance may result in reduced medical staffing during the meeting dates. We sought to examine whether there were differences in mortality after trauma among patients admitted to the hospital during, before, and after meeting dates. Using the Japan Trauma Data Bank, we analyzed in-hospital mortality in patients with traumatic injury admitted to the hospital from 2004 to 2015 during the dates of two national academic meetings - the Japanese Association for Acute Medicine (JAAM) and the Japanese Association for the Surgery of Trauma (JAST). We compared the data with that of patients admitted with trauma during identical weekdays in the weeks before and after the meetings, respectively. We used multiple logistic regression analysis to compare outcomes among the three groups. A total of 7,491 patients were included in our analyses, with 2,481, 2,492, and 2,518 patients in the during, before, and after meeting dates groups, respectively; their mortality rates were 7.3%, 8.0%, and 8.5%, respectively. After adjusting for covariates, no significant differences in in-hospital mortality were found among the three groups (adjusted odds ratio [95% CI] of the before meeting dates and after meeting dates groups; 1.18 [0.89-1.56] and 1.23 [0.93-1.63], respectively, with the during meeting dates group as the reference category). No significant differences in in-hospital mortality were found among trauma patients admitted during, before, and after the JAAM and JAST meeting dates.

## Introduction

Appropriate medical staffing is essential to provide optimal trauma care [1]. Weekend or off-hours admission has been shown to be associated with worse outcomes in patients with acute myocardial infarction (AMI), stroke, pulmonary embolism, or those who required emergency general surgery and were admitted to the intensive care unit [2–6]. This so-called “weekend effect” could possibly be explained by reduced medical staffing and resources [7, 8].

This “national meeting effect” has been examined in recent years [9–12]. Each year, many physicians attend national academic meetings and conferences to present their work, gain new knowledge, and network. Although hospitals aim to consistently deliver high quality patient care through efficient allocation of staff physicians, medical staffing during national meetings dates may be lower than that during non-meeting dates. The “national meeting effect” in Japan has been investigated; no significant differences were observed in outcomes among patients hospitalized with AMI or cardiac arrest between meeting dates and non-meeting dates [9, 10]. Interestingly, lower 30-day mortality was found among high-risk patients with AMI, cardiac arrest, and heart failure in teaching hospitals in the United States during national cardiology meeting dates [11, 12].

Although a “weekend effect” in terms of mortality has not been detected [13–15], longer emergency department stay and increased risk for missed injuries have been demonstrated for trauma patients admitted during off-hours in a community hospital setting [16]. To our knowledge, the “national meeting effect” among trauma patients has never been well elucidated. We hypothesized that hospital mortality would be higher during the meeting dates of national scientific emergency medicine and trauma surgery professional organizations than non-meeting dates and hospital mortality would be lower after the meeting dates than before the meeting dates because of reduced staffing and the positive impact of the academic meeting on high physician performance. Our study’s aim was to compare hospital mortality after trauma among patients admitted during, before, and after national meeting dates.

## Materials and methods

### Study design and data sources

This study was designed as a nationwide retrospective cohort study. We used data from the Japan Trauma Data Bank (JTDB), which was established in 2003 with the Committee for Clinical Care Evaluation of the Japanese Association for Acute Medicine and the Trauma Surgery Committee of the Japanese Association for the Surgery of Trauma (JAST). Patients with Abbreviated Injury Scale (AIS) scores of 3 or above are recorded in the database from 264 Japanese hospitals participating in trauma research and care [17]. The registry database contains patient demographics, mechanism of injury, vital signs at the scene and on arrival, admission date, AIS scores, Injury Severity Score (ISS), treatments, and survival status at discharge from hospitals. The Okayama University Hospital ethical committee approved the study (ID 1805–020). Since patient data was extracted anonymously, the requirement for informed consent was waived.

### Study sample

We obtained annual national meeting dates of two academic organizations - the Japanese Association for Acute Medicine (JAAM) and JAST - from 2004 to 2015. The during meeting dates group included patients admitted after traumatic injury during the dates of these meetings. The before and after meeting dates groups were defined as patients admitted with trauma during the same weekdays in the weeks before and after the meetings, respectively [9, 10]. The JAAM and JAST meetings are each usually held for two or three consecutive days. For example, the 2015 JAAM meeting was held from Wednesday, October 21 through Friday, October 23; the before and after meeting dates groups included patients admitted Wednesday through Friday in the weeks before and after the meeting, respectively. In this study, patients who were 16 years of age or older admitted with traumatic injury from 2004 to 2015 were enrolled. Patients in cardiac arrest at the scene or on arrival and those without age, hospital arrival date, and in-hospital mortality data were excluded.

### Outcome measures

Our primary outcome was post-trauma in-hospital mortality from all causes among patients hospitalized during, before, and after national meeting dates.

### Statistical analysis

Comparisons among the three groups were made using the chi-square test for categorical variables and analysis of variance for continuous variables. We used multiple logistic regression analysis to compare outcomes between the three groups, with the during meeting dates group as the reference category. Adjusted odds ratios (ORs) and their 95% confidence intervals (CIs) were obtained after adjusting for age (16-39 vs. 40-64 vs. 65≤); gender; mechanism of injury (blunt or others); transfer from outside hospitals; ISS (≤8 vs. 9-15 vs. 90mmHg≤ on arrival); Glasgow Coma Scale score (≤8 vs. 9-15); presence or absence of emergency surgical or hemostatic intervention (craniotomy, thoracotomy, laparotomy, or angioembolization); and type of institution (high vs. low volume centers). Because outcomes would be better at the high-volume centers, an additional analysis was conducted by dividing the patients into two groups; high volume centers (≥1,200 cases with ISS≥9 registered for 12 years) and low volume centers (<1,200 cases with ISS≥9 registered for 12 years) [18]. A subgroup analysis was also conducted, stratifying patients with or without shock and the type of national meeting. Sensitivity analysis was conducted using alternative definitions of the before and after meeting dates groups; two, three, and four weeks before and two, three, and four weeks after meeting dates, respectively, instead of one week. A two-tailed P value of <0.05 was considered statistically significant. All analyses were performed using IBM SPSS Statistics 25 (IBM SPSS, Chicago, IL, U.S.A.).

## Results

### Patient characteristics

A total of 236,698 trauma patients were registered in the JTDB during the study period. Of those, 182,877 adult trauma patients were assessed for eligibility. After 175,386 patients were excluded due to not being admitted on eligible days, 7,491 subjects were included in our analyses, with 2,481 patients in the during meeting dates group, 2,492 patients in the before meeting dates group, and 2,518 patients in the after meeting dates group (Fig. 1). Among the three groups of patients, basic characteristics including severity of trauma and life-saving surgical procedures were similar except for the age category (Table 1).

**Fig 1.**
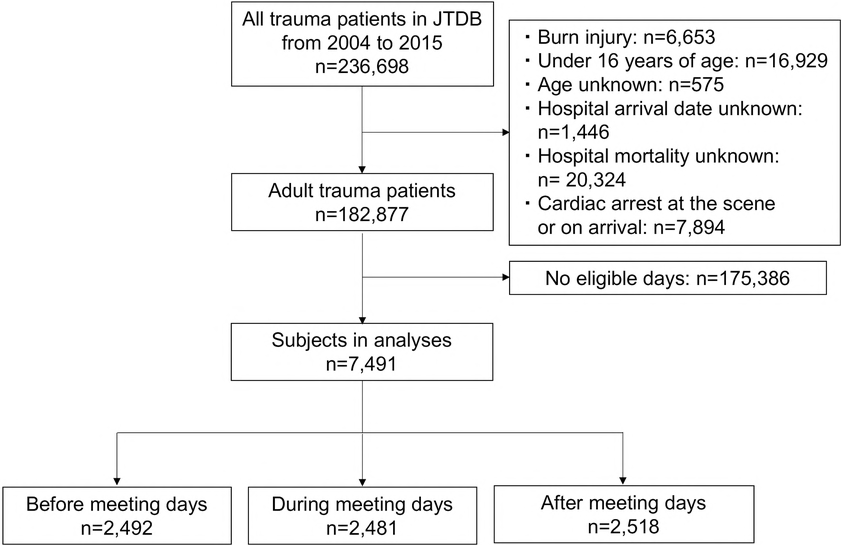
Flow diagram of the study population. JTDB, Japan Trauma Data Bank.

**Table 1.** Characteristics of trauma injury patients admitted during, before, and after national meeting dates.

|  | Before meeting<br>dates group<br>n=2,492 | During meeting<br>dates group<br>n=2,481 | After meeting<br>dates group<br>n=2,518 | P-Value |
| --- | --- | --- | --- | --- |
| Age (years) | 63 (41, 77) | 63 (41, 77) | 65 (43, 78) | 0.131 |
| 16-39, n (%) | 541 (21.7) | 604 (24.3) | 566 (22.5) |  |
| 40-64, n (%) | 750 (30.1) | 708 (28.5) | 675 (26.8) | 0.015 |
| 65≤, n (%) | 1,201 (48.2) | 1,169 (47.1) | 1,277 (50.7) |  |
| Male, n (%) | 1,557 (62.5) | 1,513 (61.1) | 1,607 (63.9) | 0.126 |
| Blunt mechanism, n (%) | 2,347 (94.2) | 2,348 (94.6) | 2,398 (95.2) | 0.249 |
| Transfers from an outside hospital, n (%) | 345 (13.8) | 367 (14.8) | 345 (13.7) | 0.485 |
| Systolic blood pressure (mmHg) | 136 (116, 158) | 136 (116, 156) | 137 (117, 158) | 0.624 |
| <90mmHg, n (%) | 180 (7.2) | 180 (7.3) | 170 (6.8) | 0.731 |
| Glasgow Coma Scale | 15 (14, 15) | 15 (14, 15) | 15 (14, 15) | 0.532 |
| ≤8, n (%) | 245 (9.8) | 242 (9.8) | 256 (10.1) | 0.874 |
| Surgical or hemostatic intervention, n (%) | 266 (10.7) | 237 (9.6) | 237 (9.4) | 0.261 |
| Craniotomy, n (%) | 101 (4.0) | 83 (3.4) | 82 (3.3) | 0.249 |
| Thoracotomy, n (%) | 32 (1.3) | 31 (1.3) | 20 (0.8) | 0.181 |
| Laparotomy, n (%) | 76 (3.1) | 72 (2.9) | 66 (2.6) | 0.652 |
| Angioembolization, n (%) | 73 (2.9) | 65 (2.6) | 78 (3.1) | 0.592 |
| Injury Severity Score | 10 (9, 20) | 10 (9, 20) | 10 (9, 21) | 0.590 |
| ≤8, n (%) | 407 (16.3) | 399 (16.1) | 398 (15.8) |  |
| 9-15, n (%) | 993 (39.8) | 980 (39.5) | 1,016 (40.3) | 0.904 |
| 16≤, n (%) | 1,009 (40.5) | 1,042 (42.0) | 1,034 (41.1) |  |
| High volume center, n (%) | 1,437 (57.7) | 1,486 (59.9) | 1450 (57.6) | 0.172 |
| Low volume center, n (%) | 1,055 (42.3) | 995 (40.1) | 1,068 (42.4) |  |
| In-hospital mortality, n (%) | 200 (8.0) | 181 (7.3) | 213 (8.5) | 0.306 |

### Comparison of mortality between the three groups

No significant differences in in-hospital mortality were observed during, before, and after meeting dates (7.3% vs. 8.0% vs. 8.5%, respectively; P=0.306; unadjusted OR [95% CI] of the before and after meeting dates groups; 1.11 [0.90-1.37], 1.17 [0.96-1.44], respectively, with the during meeting dates group as the reference category; Table 2). Even after adjusting for covariates, no significant differences in in-hospital mortality were found among the three groups (adjusted OR [95% CI] of the before and after meeting dates groups; 1.18 [0.89-1.56], 1.23 [0.93-1.63], respectively, with the during meeting dates group as the reference category; Table 2).

**Table 2.** In-hospital mortality after trauma among patients hospitalized during, before, and after national meeting dates.

|  | Before meeting<br>dates group | During meeting<br>dates group | After meeting<br>dates group | P-value |
| --- | --- | --- | --- | --- |
| Overall |  |  |  |  |
| In-hospital mortality, % (n/N) | 8.0 (200/2,492) | 7.3 (181/2,481) | 8.5 (213/2,518) | 0.306 |
| Crude OR (95% CIs) | 1.11 (0.90-1.37) | Reference | 1.17 (0.96-1.44) |  |
| Adjusted OR (95% CIs) | 1.18 (0.89-1.56) | Reference | 1.23 (0.93-1.63) |  |

Adjusted OR and their 95% CIs were obtained after adjusting for age (16-39 vs. 40-64 vs. 65≤), gender, mechanism of injury (blunt or others), transfer from an outside hospital, ISS (≤8 vs. 9-15 vs. 16≤), presence or absence of shock, Glasgow Coma Scale score (≤8 vs. 9-15), presence or absence emergency surgical or hemostatic intervention, and type of institution (high vs. low volume centers). OR: odds ratio; CI: confidence interval; ISS: Injury Severity Score

OR: odds ratio; CIs: confidence intervals; ISS: Injury Severity Score; JAAM, Japanese Association for Acute Medicine; JAST, Japanese Association for the Surgery of Trauma.

### Subgroup analysis

Although high volume centers were associated with better outcomes (6.8% overall in-hospital mortality of high volume centers vs. 9.5 % for the low volume centers; P<0.001), in-hospital mortality did not differ among the three groups according to center volume (Table 3). Additional analyses were conducted by stratifying presence or absence of shock; no differences in in-hospital mortality were found among the three groups (Table 3). Also, no significant differences according to type of national meeting were found (Table 3). The same results were obtained when considering alternative definitions of the before and after meeting dates groups (Table 4).

**Table 3.** In-hospital mortality among the three groups with stratification for high vs. low volume centers, presence or absence of shock, and type of national meeting.

|  | Before meeting<br>dates group | During meeting<br>dates group | After meeting<br>dates group | P-value |
| --- | --- | --- | --- | --- |
| High volume centers |  |  |  |  |
| In-hospital mortality, % (n/N) | 7.2 (103/1,437) | 6.0 (89/1,486) | 7.3 (106/1,450) | 0.129 |
| Crude OR (95% CI) | 1.21 (0.90-1.62) | Reference | 1.24 (0.93-1.66) |  |
| Adjusted OR (95% CI)<sup>a</sup> | 1.12 (0.77-1.64) | Reference | 1.30 (0.89-1.89) |  |
| Low volume centers |  |  |  |  |
| In-hospital mortality, % (n/N) | 9.2 (97/1,055) | 9.2 (92/995) | 10.0 (107/1,068) | 0.968 |
| Crude OR (95% CI) | 0.99 (0.74-1.34) | Reference | 1.09 (0.82-1.47) |  |
| Adjusted OR (95% CI)<sup>a</sup> | 1.25 (0.82-1.90) | Reference | 1.17 (0.77-1.78) |  |
| Systolic blood pressure ≤ 90mmHg |  |  |  |  |
| In-hospital mortality, % (n/N) | 29.4 (53/180) | 35.6 (64/180) | 34.7 (59/170) | 0.413 |
| Crude OR (95% CI) | 0.76 (0.49-1.18) | Reference | 0.96 (0.62-1.50) |  |
| Adjusted OR (95% CI)<sup>b</sup> | 1.09 (0.62-1.90) | Reference | 1.14 (0.66-1.97) |  |
| Systolic blood pressure > 90mmHg |  |  |  |  |
| In-hospital mortality, % (n/N) | 6.0 (134/2,252) | 4.5 (101/2,247) | 6.0 (138/2,293) | 0.040 |
| Crude OR (95% CI) | 1.34 (1.03-1.75) | Reference | 1.36 (1.05-1.77) |  |
| Adjusted OR (95% CI)<sup>b</sup> | 1.28 (0.93-1.74) | Reference | 1.29 (0.95-1.77) |  |
| JAAM |  |  |  |  |
| In-hospital mortality, % (n/N) | 8.3 (130/1,560) | 7.4 (112/1,513) | 9.1 (147/1,612) | 0.221 |
| Crude OR (95% CI) | 1.14 (0.87-1.48) | Reference | 1.26 (0.97-1.62) |  |
| Adjusted OR (95% CI)<sup>c</sup> | 1.28 (0.90-1.80) | Reference | 1.28 (0.91-1.80) |  |
| JAST |  |  |  |  |
| In-hospital mortality, % (n/N) | 7.5 (70/932) | 7.1 (69/968) | 7.3 (66/906) | 0.950 |
| Crude OR (95% CI) | 1.06 (0.75-1.49) | Reference | 1.02 (0.72-1.45) |  |
| Adjusted OR (95% CI)<sup>c</sup> | 1.21 (0.78-1.88) | Reference | 1.18 (0.75-1.87) |  |

OR: odds ratio; CIs: confidence intervals; ISS: Injury Severity Score; JAAM, Japanese Association for Acute Medicine; JAST, Japanese Association for the Surgery of Trauma.

a Adjusted OR and their 95% CIs were obtained after adjusting for age (16-39 vs. 40-64 vs. 65≤), gender, mechanism of injury (blunt or others), transfer from an outside hospital, ISS (≤8 vs. 9-15 vs. 16≤), presence or absence of shock, Glasgow Coma Scale score (≤8 vs. 9-15), and presence or absence of emergency surgical or hemostatic intervention.

b Adjusted OR and their 95% CIs were obtained after adjusting for age (16-39 vs. 40-64 vs. 65≤), gender, mechanism of injury (blunt or others), transfer from an outside hospital, ISS (≤8 vs. 9-15 vs. 16≤), Glasgow Coma Scale score (≤8 vs. 9-15), presence or absence of emergency surgical or hemostatic intervention, and type of institution (high vs. low volume centers).

c Adjusted OR and their 95% CIs were obtained after adjusting for age (16-39 vs. 40-64 vs. 65≤), gender, mechanism of injury (blunt or others), transfer from an outside hospital, ISS (≤8 vs. 9-15 vs. 16≤), presence or absence of shock, Glasgow Coma Scale score (≤8 vs. 9-15), presence or absence of emergency surgical or hemostatic intervention, and type of institution (high vs. low volume centers).

**Table 4.** In-hospital mortality among the three groups with alternative definitions of the before and after meeting dates groups.

|  | Before meeting<br>dates group | During meeting<br>dates group | After meeting<br>dates group | P-value |
| --- | --- | --- | --- | --- |
| $\pm 2^a$ | | | | |
| In-hospital mortality, % (n/N) | 7.1 (169/2,397) | 7.3 (181/2,481) | 7.5 (190/2,544) | 0.851 |
| Adjusted OR (95% CIs) | 1.07 (0.80-1.42) | Reference | 1.27 (0.96-1.68) |  |
| $\pm 3^b$ | | | | |
| In-hospital mortality, % (n/N) | 7.6 (193/2,546) | 7.3 (181/2,481) | 8.2 (207/2,517) | 0.450 |
| Adjusted OR (95% CIs) | 1.22 (0.92-1.61) | Reference | 1.21 (0.92-1.61) |  |
| $\pm 4^c$ | | | | |
| In-hospital mortality, % (n/N) | 7.7 (196/2,538) | 7.3 (181/2,481) | 6.6 (176/2,650) | 0.316 |
| Adjusted OR (95% CIs) | 1.23 (0.92-1.62) | Reference | 0.94 (0.71-1.26) |  |

a Two weeks before and after meeting dates as the before meeting dates group and after meeting dates group.

b Three weeks before and after meeting dates as the before meeting dates group and after meeting dates group.

c Four weeks before and after meeting dates as the before meeting dates group and after meeting dates group.

Adjusted OR and their 95% CIs were obtained after adjusting for age (16-39 vs. 40-64 vs. 65≤), gender, mechanism of injury (blunt or others), transfer from an outside hospital, ISS (≤8 vs. 9-15 vs. 16≤), presence or absence of shock, Glasgow Coma Scale score (≤8 vs. 9-15), presence or absence of emergency surgical or hemostatic intervention, and type of institution (high vs. low volume centers).

OR: odds ratio; CIs: confidence intervals; ISS: Injury Severity Score.

## Discussion

In this study, we investigated whether there was a difference in mortality among patients admitted due to traumatic injuries during, before, and after dates of national academic acute medicine and trauma meetings. Contrary to our hypothesis, we found no significant differences in in-hospital mortality among the three groups, even after adjusting for measurable confounders.

To our knowledge, “national meeting effects” were first investigated in the United States, focusing on national cardiology meetings; lower staffing and differences in composition of physicians during the meeting dates were found to possibly affect treatment utilization and outcomes [11]. In this study, no significant differences in mortality of AMI patients were found between those in the hospital during meeting and non-meeting dates; however, high-risk patients with AMI, cardiac arrest, and heart failure admitted to teaching hospitals during meeting dates were found to have lower mortality than those admitted during non-meeting dates [11]. The present study is the first to examine the “national meeting effect” regarding mortality in trauma patients.

Previous studies have not detected the “weekend effect”; admission on nights or weekends for trauma patients was not associated with increased mortality [13–16, 19, 20] or even better outcomes [21]. Generally, a plausible explanation for the “weekend effect” includes several factors such as reduced medical staffing, decreased access to some tests and procedures, and the influence of variations in case mix [19, 20, 22]. For trauma patients, hospitals are explicitly required to be appropriately staffed and to provide optimal care, regardless of when injured patients are admitted [21, 23]. However, trauma patients presenting off-hours were more likely to have missed injuries, [16, 24], in particular thoracic spine or abdominal injuries [24]. Specifically, Schwartz DA, et al. showed that off-hour presentation of pelvic fracture patients with hemorrhagic shock caused a delay in door-to-angioembolization time, resulting in increased mortality [23].

In theory, variations of patient characteristics would not be influenced by admission dates occurring during, before, and after national meetings. While relatively less experienced emergency physicians or trauma surgeons providing leadership would be expected during academic national meeting dates, strong leadership, teamwork, and technical skills are essential components for team performance and patient care in initial trauma management [25, 26]. Our results showed no significant differences in mortality among those admitted during, before, and after national meeting dates; these findings may be explained by the possibility that every hospital strives to consistently deliver high quality care. Although no clear evidence exists that national academic meetings directly improve medical staff performance and positively impact patient outcomes, hospital participation in a trauma quality improvement program has been demonstrated to be associated with better patient outcomes [27]. Hence, we compared hospital mortality among patients admitted during, before, and after national meeting dates to investigate the “post-national meeting effect,” assuming that participation in national academic meetings has a positive effect on clinical performance and patient outcomes. Despite our hypothesis, our results detected no “post-national meeting effect.”

Our study has several limitations. First, we could not identify differences in medical staffing among the three groups, such as composition of emergency physicians vs. trauma surgeons who treated the patients; therefore, we are unable to explain why no significant differences in mortality were found. Second, the influence of national meeting geographical regions and locations was not accounted for, which may have affected our results. Third, as Jena AB, et al. showed that high-risk AMI patients admitted to teaching hospitals during national meeting dates received less percutaneous coronary intervention, hospital type should have been taken into consideration [11]. We performed additional analyses according to hospital volume, in which the same results were obtained. Fourth, possible confounders, including comorbidities, were unavailable in this study. Fifth, although major tertiary hospitals providing high-quality trauma care participate in the JTDB, a degree of random error and selection bias may have occurred, as this was not a comprehensive study [17]. Finally, since we focused on national academic meetings only in Japan, our results could not be applied to other countries, considering differences in the settings and geography of Japanese healthcare systems.

## Conclusions

We observed no significant differences in in-hospital mortality after trauma among patients admitted during, before, and after national acute medicine and trauma meeting dates. As hospitals are assumed to be struggling to consistently provide optimal care for trauma patients, participating in these meetings is acceptable for sharing and generating new knowledge. Further population-based studies are required to validate our results.

## Acknowledgements

We express our gratitude to all the participants in the JTDB registry, members of the Trauma Registry Committee (Japanese Association for Trauma Surgery), and the Committee for Clinical Care Evaluation (Japanese Association for Acute Medicine).

## Supporting information captions

Not applicable.

